# Protein aggregation inhibitors induce divergent transcriptional responses in a cellular model of ɑ-synuclein seeded aggregation

**DOI:** 10.64898/2026.01.29.701691

**Authors:** Paulien Van Minsel, Chris Van den Haute, Eline Vonck, Shirine Hentati, Michele Curcio, Xinran Song, Qian Yu, Matthias Versele, Kenneth W. Young, Patrick Chaltin, Bernard Thienpont, Veronique Daniels, Veerle Baekelandt, Wouter Peelaerts

## Abstract

Parkinson’s disease (PD), dementia with Lewy Bodies (DLB) and multiple system atrophy (MSA) are progressive neurodegenerative disorders marked by the pathological aggregation of alpha-synuclein (ɑSyn). Despite significant research efforts, effective therapeutic interventions remain elusive due to limited understanding of the cellular effects of ɑSyn aggregation and propagation. This study presents the development of a scalable cellular seeding assay for screening small molecules targeting cellular ɑSyn seeded aggregation. By leveraging a fluorescent reporter of ɑSyn and phenotypic screening, the assay enables high-throughput evaluation of potential inhibitors in a cellular environment mimicking disease pathology. We evaluated three different αSyn aggregation inhibitors tested in clinical trials for PD: Minzasolmin, Emrusolmin and EGCG and profiled gene expression using multiplexed single cell RNA sequencing in order to examine their distinct effects on cellular pathways associated with ɑSyn overexpression or seeded aggregation. Two cellular activities were prominently affected: lipid metabolism and rRNA processing. Notably, while EGCG effects were confined to cells with aggregated αSyn, Minzasolmin and Emrusolmin also produced transcriptional changes in cells without aggregated αSyn. Each of the compounds tested induced a partial reversal of transcriptional effects resulting from αSyn seeded aggregation. We identified 391 genes that were no longer significantly differentially expressed upon addition of compound, relative to cells with seeded aggregation. This platform bridges phenotypic screening and molecular pathway analysis, providing insights into druggable pathways for synucleinopathies. The molecular signatures identified here can assist in testing and benchmarking future drug discovery leads.

## Introduction

Parkinson’s disease (PD), dementia with Lewy Bodies (DLB) and multiple system atrophy (MSA) are progressive neurodegenerative disorders characterized by the pathological aggregation of ɑ-synuclein (ɑSyn). Despite extensive research, the exact mechanisms underlying cellular ɑSyn aggregation and its propagation remain poorly understood, limiting effective therapeutic development. The aggregation and subsequent transmission of ɑSyn fibrils are recognized as critical factors contributing to PD progression^1,2^, making the discovery of small molecules capable of inhibiting or modulating this process a top priority in therapeutic research. Development of compounds that inhibit αSyn aggregation or prevent its transmission represents a rational therapeutic strategy for preventing PD onset or progression

Although the precise reasons for ɑSyn aggregation in its cellular environment remain unclear, this process is significantly influenced by ɑSyn protein expression levels, its interactions with lipids, and external triggers that can cause ɑSyn protein modifications or acutely increase ɑSyn expression^3^. The misfolding of ɑSyn monomers into stable fibrils requires multiple steps, beginning with its initial misfolding into unstable, amorphous oligomers and finally forming a stable fibrillar nucleus that can seed the formation of new fibrillar aggregates^4,5^. These fibrillar aggregates can form large inclusions, or Lewy bodies, which, as they mature, sequester lipids, organelles and other biomolecules^6^ that prevent the cell from functioning properly, leading to cellular damage and cell death^7,8^.

Physiologically, αSyn is important for regulating synaptic and immune function in healthy cells. Compounds that are being developed for PD, DLB or MSA therapy therefore require selectivity towards aggregated αSyn to ensure treatment safety. Furthermore, given the interactions of aggregated ɑSyn within its cellular environment it is important to implement the relevant tools to study and elucidate druggable pathways via which ɑSyn aggregation can be targeted. Current therapeutic strategies target different aspects of αSyn pathology, either preventing aggregation of misfolded protein, enhancing the clearance of the aggregates, or inhibiting their transmission.^9^ Recent advances in cellular modeling and high-throughput screening have identified various small molecule candidates that show promise in modulating ɑSyn aggregation, preventing inclusion formation and improving cellular health^2^. Previous studies have identified several compounds that can inhibit α-syn aggregation, including Minzasolmin (UCB0599), Emrusolmin (Anle138b), epigallocatechin-3-gallate (EGCG), NPT100-18A and SynuClean-D.^10^ Some of these, i.e. Minzasolmin, Emrusolmin and EGCG, have undergone clinical trials but the cellular pathways they affect remain incompletely clear, which represent the focus of our work.

Advances in single-cell RNA sequencing (scRNA-seq) over the past decade enabled the simultaneous profiling of multiple compound treatments in a single screen, significantly reducing the associated costs and bypassing batch effects. Here, we present a scalable cellular aggregation assay and a phenotypic screening model that focuses on ɑSyn aggregation via seeding with pre-formed fibrils (PFFs). We use a multiplexed scRNA-seq approach to investigate how three previously identified ɑSyn aggregation inhibitors alter the transcriptome of neuroglioma cells. We characterize gene expression changes induced by acute αSn aggregation neurotoxicity, and the counteracting activity of ɑSyn modulators. Our findings thus outline the therapeutic potential of these compounds and deliver a strategy to define and select interesting novel hits in future studies.

## Materials and Methods

### Viral vector production

Lentiviral vectors were produced as described previously^11^. In brief, HIV-derived lentiviral vectors pseudotyped with the VSV-G envelope were produced by transient transfection of HEK 293T cells. The transfer plasmid included human wild type αSyn or αSyn-YFP as transgenes under control of a constitutive CMV promoter, with the WPRE to boost transgene expression, and the cPPT/CTS for efficient nuclear import. For the attachment of YFP as a reporter protein, a short but flexible linker is used at the C-terminus of αSyn. To allow selection of transduced cells, an IRES-puromycin selection cassette was introduced. Following triple transient transfection, secreted vector particles were harvested from the HEK 293T cell medium, purified through filtration and concentrated through centrifugation using Amicon Ultra-15 centrifugal filters (100K, Millipore). p24 antigen content was determined using the HIV-1 p24 Core Profile ELISA and vectors were stored at −80°C for further use.

### Generation and sonication of pre-formed fibrils

Recombinant monomers of human αSyn were obtained from Proteos (MJFF). Upon arrival, monomers were stored at −80°C. Generation of PFFs from these monomers was done according to the MJFF protocol. In short, aSyn monomers were thawed at room temperature and diluted in sterile PBS at a concentration of 5 µg/µl. Next, they underwent controlled *in vitro* aggregation at 37°C at a speed of 1000 rpm in an Eppendorf® ThermoMixer® for 7 days. After aggregation, samples are aliquoted and stored at −80°C until the start of the experiment. Right before use, 10 µl fibrils were thawed at room temperature, dissolved in 200µl OptiMEM and immediately sonicated in a waterbath (VWR, ultrasonic cleaner) at 4°C during 20 minutes. The PFFs were characterized by negative uranyl acetate staining followed by TEM to measure the length. The average length measured for αSyn PFFs after waterbath sonication was 83 nm ± 52 nm (mean ± SD, 1000 measured particles).

### Generation of cell lines and seeding assay

Human H4 neuroglioma cells were transduced with lentiviral vectors expressing human αSyn or human αSyn-YFP by adding the viral vector to the cell medium for 24 hours. Cells were subsequently treated with 2 µg/mL puromycin to select stably expressing cells. Stably transduced cells were frozen in liquid nitrogen until further use.

### Seeding assay for compounds analysis

αSyn or αSyn-YFP expressing cells were seeded at a density of 6,500 cells in 25 µl in a 384 well plate. After overnight culture cells were incubated with recombinant sonicated αSyn pre-formed fibrils at a concentration of 2.5 µg/mL. To facilitate the entry and uptake of the PFFs in the cell culture, sonicated PFFs were mixed with lipofectamine 2000 to allow complexation in a ratio of 5 µg PFFs/20 µl lipofectamine for 30 minutes in OptiMEM medium. Fifty µl of this PFF/lipofectamine complex was added to the cells. After 3 hours of incubation 25 µl of 4-fold concentrated compound was added to the cells for 24 hours. Finally, cells were washed with PBS, fixed with 4% paraformaldehyde and stored in PBS with DAPI (200 ng/ml) for further analysis.

### Seeding assay for scRNA-seq

αSyn expressing cells, were seeded at a density of 25,000 cells in 50 µl in a 96 well plate. After overnight culture, the cells were incubated with a complex of lipofectamine and recombinant αSyn pre-formed fibrils at a concentration of 2.5 µg/mL. Fifty µl of lipofectamine/PFF complex was added to the cells resulting in a final concentration of 1.25 µg/ml PFFs. After 2 hours of incubation with PFFs 25 µl of 5-fold concentrated compound was added to the cells. DMSO-only condition was used as a negative control. After 24 hours of incubation with compounds, cell labeling was performed using the lipid barcode-based MULTI-seq technique as previously described by McGinnis *et al*.^12^ with minor modifications, using MULTI-seq Lipid Modified Oligos (Sigma-Aldrich, cat. no. LMO001). We used Capture Sequence 1 compatible with the Feature Barcoding technology of 10x Genomics for capturing the barcode fraction. Cells were dissociated in a TrypLE (Thermo Fischer, A1285901) solution with 20 nM of the equimolar Anchor:Barcode mix. After 5 mins at 37°C, the co-anchor reagent was added at a final concentration of 20 nM and cells were incubated for another 5 min at 37°C. Samples were washed in the 96-well plate with ice-cold 1% BSA (Thermo Fisher, 15260037) in DPBS (Thermo Fisher, A1285901) for a total of three times. Labelled cells were then pooled together in a 50 ml tube prefilled with 30 ml 1% BSA in DPBS. Following another two washes with 1% BSA, cells were resuspended in 0.04% BSA in DPBS to reach 1000 cells/µl and run through a 40 µM Flowmi cell strainer (Sigma Aldrich, BAH136800040) to obtain single cells. Finally, cells were loaded on the Chromium single cell platform (10x Genomics) and continued with library preparation.

### Immunocytochemistry

For antigen detection of αSyn aggregates, primary antibodies listed in **Table 1** were used at 4°C for overnights staining. Slides were triple washed in 0.1% Triton-X in PBS or PBS only and incubated with secondary antibody (**Table 1**) and DAPI (1:1000 – 1:3000) for two hours at room temperature after which the sections were washed again with 0.1% Triton-X in PBS or PBS. To visualize αSyn aggregates we further incubated the sections for 30 minutes with Amytracker 680 (Ebba Biotech) diluted at concentration of 1:5000 in PBS. The slides were triple washed in PBS before being sealed with Vectashield Antifade Mounting Medium (Vector Laboratories) or Mowviol Mounting Medium.

**Table 1.** Antibodies used in study for immunocytochemistry.

| Antibody | Concentration | Application | Supplier | Catalog nr. (clone) |
| --- | --- | --- | --- | --- |
| Rabbit anti-α-synuclein | 1:1000 | IF | Abcam | Ab138501 (MJFR-1) |
| Rabbit anti-PSer-129 α-synuclein | 1:500-1000 | IHC/IF | Abcam | Ab51253 (EP1536Y) |
| Mouse anti-PSer-129 α-synuclein | 1:500 | IF | Abcam | Ab184674 (81a) |
| Goat anti-GFP | 1:1000 | IF | Abcam | Ab6673 |
| Rabbit anti-P62 | 1:500 | IF | Proteintech | 55274-1-AP |
| Amytracker 680 | 1:5000 | IF | EbbaBiotech |  |
| Donkey anti-rabbit Alexa-488 | 1:1000 | IF | ThermoFisher | A-21206 |
| Donkey anti-goat Alexa-488 | 1:500 | IF | Invitrogen | A11055 |
| Donkey anti-rabbit Alexa-555 | 1:1000 | IF | ThermoFisher | A-31572 |
| Donkey anti-mouse Alexa-555 | 1:500 | IF | Invitrogen | A-31570 |
| Donkey anti-rabbit Alexa-647 | 1:500-1000 | IF | ThermoFisher | A-31573 |

### Cellular phenotyping

For the analysis of αSyn-YFP cells, no additional staining was performed, and plates were directly imaged using the Operetta High Content Imaging System (PerkinElmer) or the CellInsight CX7 High content Screening platform (Thermo Scientific). Up to 6 images are taken with one well, at 20X resolution to visualize a number of cells. For analysis of Pser129-αSyn, additional staining was performed before analysis. The number of αSyn-YFP aggregates and Pser129-αSyn-positive cells was quantified using YFP and via immunofluorescent staining against pSer129-αSyn, respectively. One count is identified as a cellular inclusion after cellular segmentation. The total number of cell counts was determined via DAPI staining. To assess cellular toxicity, the ratio of treated versus non-treated DAPI-positive cells is given as a % of surviving cells. Analysis was done via the Thermo Scientific HCS Studio Cell Analysis Client Software.

### scRNA-seq

scRNA-seq libraries were prepared according to the Chromium Single Cell 3’ Reagent Kits v3.1 with Feature Barcoding Technology User Guide (CG000389, 10x Genomics). For the generation of the barcode specific library the *MULTI*-seq additive primer (5′-CCTTGGCACCCGAGAATTCC-3′) was added during cDNA amplification. After which, barcode and endogenous cDNA fractions were separated using SPRI size selection, allowing for the construction of an independent MULTI-seq barcode and a gene expression library. For the endogenous transcript fraction, the protocol was continued according to the user guide. For the MULTI-seq barcode fraction, an additional 2x SPRI clean-up was performed. Lastly, an indexing PCR was performed to add Illumina adapters and sample specific indexes using the Feature SI Primer 3 (10x Genomics, PN2000263) and TruSeq RPIX primer. After library quality control using a Bioanalyzer high sensitivity DNA analysis kit (Agilent, 5067-4626), libraries were sequenced on an Illumina NovaSeq6000 and an Illumina NextSeq2000 instrument, with 28 and 91 base-pairs for read 1 and 2, respectively.

### scRNA-seq data processing

Single-cell RNA-seq data were processed and analyzed using the Seurat (v5.3.0)^13^ R package. Cell cycle phase was assigned by applying *CellCycleScoring* function. Feature selection was carried out using *FindVariableFeatures*, where the top 2,000 highly variable genes were identified for downstream analysis. Principal component analysis (PCA) was subsequently conducted with the *RunPCA* function on the selected variable features. The top 20 principal components were then used for neighborhood graph construction by *FindNeighbors* function. Dimensionality reduction for visualization was performed with Uniform Manifold Approximation and Projection (UMAP) via the *RunUMAP* function. Louvain algorithm was employed for clustering using *FindClusters* with parameter “resolution = 1”.

Demultiplex2 (v1.0.1)^14^ package was used to assign sample barcode to cells, only cells with confident single barcode assigned were retained for subsequent analysis. After filtering for mitochondrial percentage (<15%) and RNA count (1,500 – 20,000), we retained 2,652 cells with 6,795 mean *reads* per cell.

### Differential expression analysis

Differential expression analysis was performed using the R/edgeR package (v4.6.2)^15^ on raw count matrices extracted from Seurat objects. For each comparison, cells were subsetted based on metadata annotations, and raw counts were modeled using a quasi-likelihood negative binomial generalized log-*linear* model by calling glmQLFit() and glmQLFTest() functions. To account for technical variation, cellular detection rate (CDR) was included as a covariate in the design matrix. Genes were considered significantly differentially expressed if they met the following criteria: false discovery rate (FDR) < 0.05, absolute log₂ fold change > 0.26 (corresponding to a 20% change in gene expression), and expression in >10% of cells in either group.

Volcano plots were generated using R/ggplot2 (v3.5.2), with gene-wise log₂ fold change plotted against –log₁₀(*p*-value or FDR). Genes passing significance thresholds were color-coded to indicate directionality of regulation (upregulated or downregulated). Density-based coloring was applied to highlight point density across the plot. Top 10 up- and down-regulated genes were labeled based on ranking by fold change. For the pathological condition, PD-associated genes retrieved from the Open Targets Platform were highlighted using green triangles.

### Gene Set Enrichment and GO analysis

GO enrichment was performed using the clusterProfiler package (v4.16.0)^16^ in R. Differentially expressed genes were filtered based on adjusted p-value (< 0.05), absolute log₂ fold change (> 0.26), and expression in >10% of cells in either group. GO over-representation analysis (ORA) was conducted using the enrichGO() function with the org.Hs.eg.db (v 3.21.0) annotation database, focusing on the Biological Process (BP) ontology. The background gene universe consisted of all genes tested, ranked by log₂ fold change.

In parallel, GSEA was performed using the gseGO() function, with genes ranked by log₂ fold change across the full dataset. Gene sets were filtered to include those with sizes between 100 and 500 genes. Both analyses used the Benjamini-Hochberg method for multiple testing correction, with significance thresholds set at adjusted p-value < 0.05. Visualization of enriched terms was performed using dot plots, highlighting the top 10 categories based on adjusted p-value and gene count.

For selected gene subsets, including PD-associated genes and those reversed by compound treatment, GO enrichment analysis was performed using the enrichGO() function from the R/clusterProfiler package (v4.16.0). Analyses were conducted separately for the BP and Molecular Function (MF) ontologies using the org.Hs.eg.db annotation database. Enrichment was assessed using the Benjamini-Hochberg method for multiple testing correction (p-adjusted < 0.05).

These targeted analyses were used to identify functional themes within curated gene sets, such as those implicated in αSyn–mediated pathology or transcriptional reversal following compound treatment. Results were visualized by barplot, size determined by gene count and color by FDR adjusted p-value.

## Results

### Cellular seeding assay

To develop a phenotypic screening assay for compound testing and functional transcriptomics we developed a scalable cellular ɑSyn-seeded aggregation assays. The cellular seeding assay is based on the seeded aggregation of human αSyn with human PFFs in human αSyn overexpressing H4 neuroglioma cells. In the overexpressing cells, αSyn was fused to YFP via a flexible but covalent linker to directly visualize the expression, seeding and aggregation of αSyn. To generate a stably expressing αSyn-YFP line, cells were transduced with LV CMV-αSyn-YFP-Puro viral vectors (**Fig. 1a**). By addition of puromycin transduced cells were selected. The resulting cell population exhibited stable expression of αSyn-YFP, which is visualized as fluorescent diffuse YFP staining (**Fig. 1b**).

**Figure 1.**
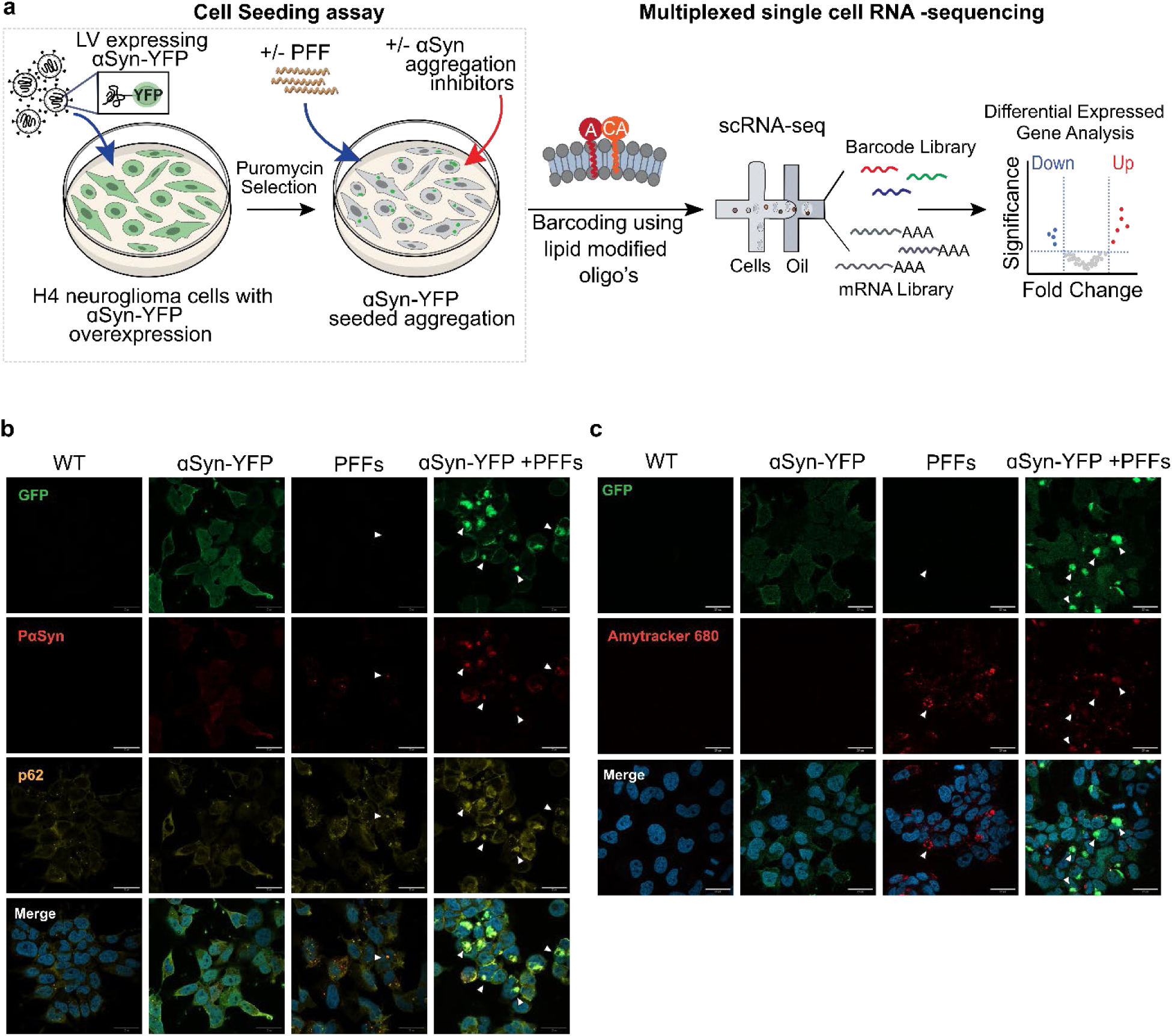
Cellular seeding with PFFs causes ɑSyn aggregation. ***(a)*** H4 neuroglioma cells overexpressing ɑSyn-YFP are treated with PFFs to induce seeded aggregation, transitioning from diffuse expression to discrete ɑSyn-YFP inclusions. Subsequently, cells are labelled using lipid modified oligo’s and pooled for single cell RNA sequencing. ***(b)*** Representative images showing ɑSyn-YFP fluorescence in untreated cells (left) and cells treated with PFFs (right). Diffuse fluorescence is replaced by discrete puncta indicative of aggregation. Nuclei are stained with DAPI (blue). Scale bars: 50 µm (top) and 20 µm (bottom). ***(c)*** High-magnification images showing co-localization of ɑSyn-YFP inclusions (YFP, green) with pɑSyn-positive aggregates (yellow, arrowheads), confirming their pathological nature. Scale bar: 10 µm.

To initiate seeded aggregation of αSyn, PFFs were introduced to the αSyn-YFP-expressing H4 cells via transfection with lipofectamine. The fibrils used for this assay are assembled in PBS shaking at 37°C under standard conditions. The fibrils were sonicated in a cooled water bath sonicator, to yield PFFs with an average length of 83 nm ± 52 nm (mean ± SD) (**Fig. S1**). Before addition to the cell medium, a complex with PFFs and lipofectamine was formed to facilitate the uptake of PFFs into H4 cells. The concentration of PFFs added to the cell culture is 2,5 µg/mL. The PFFs act as seeds, promoting the aggregation of the overexpressed αSyn-YFP. As a result, the initial diffuse staining of αSyn-YFP became dense and localized in the cytoplasm. Fluorescent images show the formation of αSyn-YFP aggregates, indicated by bright fluorescent puncta (**Fig. 1b**).

This happened within several hours, with puncta forming already after 6 hours. The seeding reaction with αSyn-YFP reached a plateau at 24 hours (**Fig. S2b**). These inclusions were further characterized by positive staining for the pathological marker PSer129-αSyn (**Fig. 1b**), that also colocalized with p62, a biochemical signature of pathological synuclein aggregates. In addition, the PSer129-αSyn+ aggregates were also found to be positive for Amytracker, a molecular probe that specifically binds with β-sheet amyloids (**Fig. 1c**). The colocalization of αSyn-YFP with PSer129-αSyn and Amytracker highlights the conversion of αSyn-YFP into amyloid-like aggregates. It also indicates successful seeding and aggregation of αSyn-YFP in response to the exogenously added PFFs.

To confirm that exogenous fibrils can also seed αSyn aggregation in H4 αSyn overexpressing cells without a YFP tag, we repeated the cellular seeding assay with untagged αSyn. Results show that upon addition of PFFs, cellular inclusions formed, and that these inclusions were positive for PSer129-αSyn (**Fig. S2c**).

To efficiently characterize the seeded aggregation of αSyn, automated cellular segmentation and inclusions counts were performed during fluorescent scanning to quantify the number of inclusions per cell. Only cells that had a bright and punctuate inclusion of αSyn-YFP were counted as a positive inclusion. DAPI was used to count the total number of cells and was used as a measure of cellular health. A decrease in total number of cells versus a non-treated condition would be indicative of cell loss and cellular toxicity. Upon treatment of PFFs we did not observe a loss in total number of cells, indicating that within 24 hours the seeded aggregation of αSyn results in detectable cellular inclusions but in the absence of significant cell loss (data not shown)

### Single cell transcriptional profiling of an α-syn aggregation model

As our seeding model efficiently reproduced αSyn pathology *in cellullo*, we used this model to understand the effect of αSyn aggregation inhibitors on gene expression in the context of αSyn cellular toxicity. We selected three well characterized compounds, epigallocatechin gallate (EGCG), Minzasolmin (UCB0599) and Emrusolmin (Anle-138), which are known to interact with αSyn multimers, αSyn aggregates in solution or at the membrane interface with αSyn^17–20^. EGCG is a natural antioxidant found primarily in green tea that has been extensively studied in PD, MSA and other neurodegenerative disorders^21^. It has potential effects on αSyn aggregation, via direct binding, but also on oxidative and inflammatory pathways^21^. Emrusolmin or Anle138b is a small molecule developed to target aggregated prion proteins and aggregated αSyn^22^. It has been tested in preclinical and early phase clinical studies of PD^23^. Minzasolmin is a small molecule that binds with αSyn aggregates^24^.

Specifically, the three compounds (Minzasolmin, Emrusolmin and EGCG) were added 3 hours after cellular seeding was initiated with PFF (three hours after addition of the PFF and lipofectamine complexes to the cell medium). This is to ensure that the compounds interact with pathways relevant to αSyn seeding and not with the uptake of PFF complexes. We first determined their efficacy and the working concentrations in the cellular αSyn seeding assay. All three compounds effectively inhibited αSyn seeded aggregation albeit at varying efficiencies (**Fig. 2a**). EGCG was most potent with an IC_50_ of 0,51µM, whereas Minzasolmin, Emrusolmin had higher IC_50_ values of 12,63µM and 20,98 µM, respectively. Next to testing the inhibitory effects of the compounds with the αSyn-YFP overexpressing line, we further tested the effects of the same compound in a cell line overexpressing untagged αSyn. Here, we observe that EGCG was again most potent with an IC_50_ of 0,27µM, whereas Minzasolmin and Emrusolmin had higher IC_50_ values of 19,84µM and 34,97 µM, respectively (**Fig. 2b**). For both conditions (αSyn-YFP and αSyn overexpression) all three inhibitors had comparable dose response effects, indicating that the YFP-tag does not interfere with the inhibition of seeded αSyn aggregation. To control for cellular toxicity, nuclear counts were performed. After treatment with EGCG, no cellular toxicity was observed at the highest tested dose (**Fig. 2c**). However, higher doses of Minzasolmin and Emrusolmin led to cellular toxicity.

**Figure 2.**
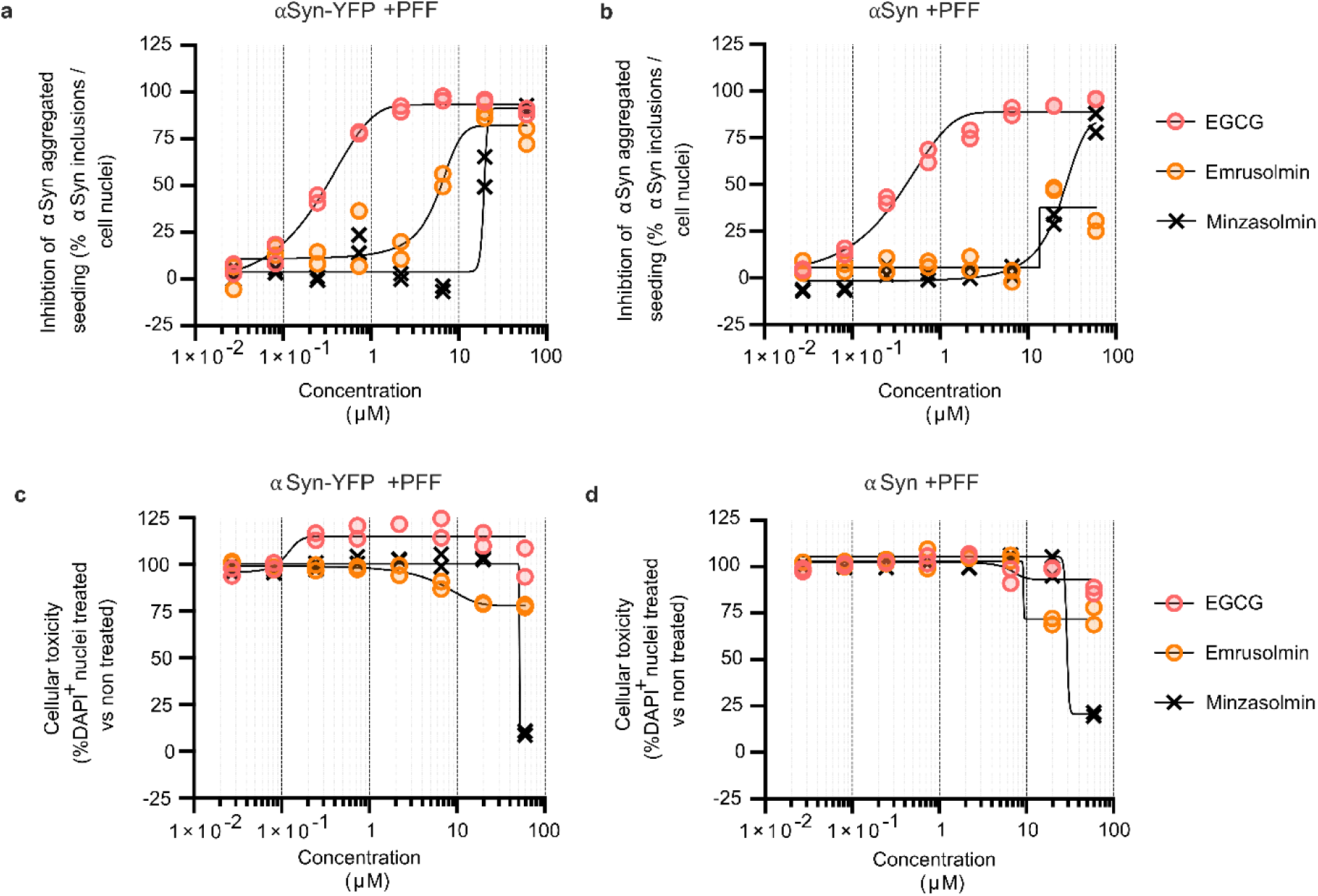
Dose response curve of EGCG, Emrusolmin and Minzasolmin on αSyn seeded aggregation and compound toxicity. ***(a)*** Concentration-dependent inhibition of αSyn-YFP aggregation in the cellular seeding assay measured by fluorescent αSyn-YFP inclusions. ***(b)*** Concentration-dependent inhibition of αSyn aggregation in the cellular seeding assay measured by fluorescent pSer129-αSyn fluorescent staining of αSyn inclusions. ***(c)*** Cellular toxicity assessed by nuclear counts in αSyn-YFP and **(d)** αSyn seeded conditions with PFFs following compound exposure. Data are expressed relative to DMSO treated condition. Shown are representative experiments with two dose response series (n = 2).

We next tested the transcriptomic effects of the three aggregation inhibitors on αSyn-YFP overexpressing cells with or without seeded PFFs. We included DMSO-only and CAY10603 inhibitor conditions as negative and positive control, respectively. CAY10603 is a potent HDAC6 and to a lesser extent HDAC1/2/3 inhibitor^25^. Cells were treated with the compounds for 24 hours after which they were labeled per treatment modality using a separate oligonucleotide barcode according to the MULTI-seq workflow^26^. To avoid batch effects, all 12 treatment combinations were assessed in a single multiplexed experiment. Following sequencing, we obtained an average of 221 cells per treatment passing quality filtering, in which we detected a mean of 10,269 unique transcripts and 3,702 unique genes per cell (**Fig. S3a, Table 2**). Cells were assigned to conditions using the deMULTIplex2 classification algorithm (**Fig. S3b**)^14^. Dimensionality reduction revealed distinct transcriptional profiles across treatment conditions, indicating that each compound induces unique gene expression signatures (**Fig. S4**). As anticipated, the positive control CAY10603 induced a highly distinct transcriptional profile, characterized by, among others, upregulation of *AREG* and *TFPI-2* (**Fig. S4d**) ^27,28^. This thus validates our lipid barcoding multiplexing workflow.

**Table 2.** Summary quality control metrics of single cell RNA data.

| Condition | Number of cells | Mean unique transcripts per cell | Median transcripts per cell | Mean unique genes per cell | Median unique genes per cell |
| --- | --- | --- | --- | --- | --- |
| Control, DMSO | 493 | 10272 | 9624 | 3723 | 3653 |
| Control, DMSO +PFF | 326 | 10577 | 10102 | 3772 | 3774 |
| CAY10603, high | 107 | 10863 | 10287 | 3760 | 3712 |
| CAY10603, high + PFF | 161 | 10790 | 10247 | 3730 | 3708 |
| CAY10603, low | 130 | 10726 | 10344 | 3744 | 3673 |
| CAY10603, low + PFF | 174 | 11044 | 10982 | 3770 | 3802 |
| Minzasolmin | 282 | 9834 | 9436 | 3607 | 3548 |
| Minzasolmin +PFF | 151 | 10312 | 10089 | 3725 | 3687 |
| Emrusolmin | 187 | 9560 | 8995 | 3588 | 3420 |
| Emrusolmin +PFF | 204 | 9823 | 9296 | 3629 | 3623 |
| EGCG | 272 | 9934 | 9558 | 3667 | 3650 |
| EGCG +PFF | 165 | 10190 | 9547 | 3751 | 3762 |
| **All** | **2652** | **10269** | **9817** | **3702** | **3670** |

### Compounds modulating α-syn aggregation produce divergent transcriptional effects

We first assessed the effect of α-syn modulators on the H4 transcriptome in the absence of αSyn PFFs, to disclose the overall transcriptomic impact of each compound. Treatment with Minzasolmin, Emrusolmin, and EGCG resulted in 134, 598 and 9 differentially expressed genes respectively (DEGs) (**Fig. 3a-c**), with little to no overlap between DEG sets (**Fig. 3e**). Minzasolmin treatment reduced expression of several interferon-induced genes, e.g. *IFIT1* and *IFIT3*, while inducing expression of genes involved in sterol metabolism, e.g. *FDPS* and *MSMO1*. Interestingly, several processes associated with viral response were suppressed after treatment with Minzasolmin (**Fig. 3d**). Emrusolmin primarily reduced expression of mRNA processing and splicing-related genes, *e.g. TENT5A*, and induced expression of lipid and xenobiotics metabolism-related genes, *e.g. MT2A*. Genes which showed differential expression in response to EGCG relate to the tumor necrosis factor (TNF) pathway, *e.g. IRF1* and *BIRC2*. No genes were commonly regulated by all three inhibitors; however, Minzasolmin and Emrusolmin shared 48 differentially expressed genes, corresponding to 57% of all DEG in Minzasolmin, suggesting partial overlap in their transcriptional impact (**Fig. 3e**). Some of these effects align with their reported mechanism of action. Indeed, Emrusolmin interferes with aggregate formation, and downregulates pathways promoting protein expression as well as mRNA processing, suggesting an attenuated protein production.^29^ Finally, EGCG affected TNF pathway genes, in line with previous reports indicating EGCG’s role in modulating TNF-α signaling to attenuate inflammation (**Fig. 3c**).^30,31^ However GSEA showed no pathways significantly activated or suppressed in the EGCG condition, likely due to the limited number of genes differentially expressed. In conclusion, each of these compounds thus had a disparate effect on gene expression which could in part be explained by their reported mechanisms of action.

**Figure 3.**
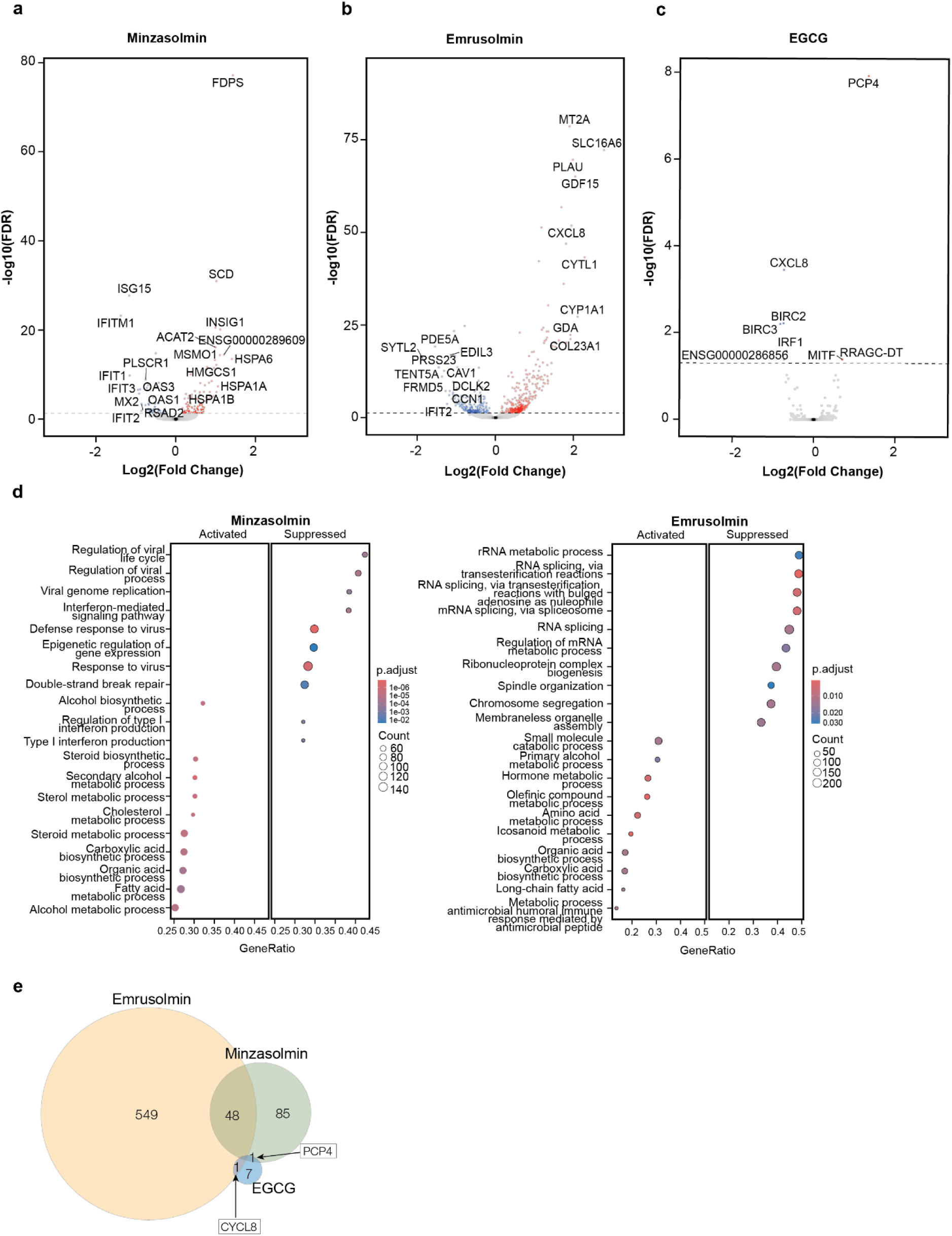
Distinct transcriptional activation by α-synuclein aggregation inhibitors in cells highlights compound-specific pathogenic signatures. ***(a-c)*** Volcano plots showing differential gene expression between cells treated with Minzasolmin (a), Emrusolmin (b), or EGCG (c) versus DMSO-treated controls under non-PFF conditions. Each dot represents a gene; red and blue indicate significantly upregulated and downregulated genes, respectively. Dashed lines denote FDR = 0.05. ***(d)*** Dot plots display gene set enrichment analysis (GSEA) results based on all genes ranked by log₂ fold change, highlighting pathways enriched among upregulated and downregulated genes. Results for Minzasolmin and Emrusolmin. Pathways are filtered by gene set size (100–500 genes) and significance threshold (padj < 0.05). Top 10 upregulated and 10 downregulated pathways are shown. **(e)** Venn diagram illustrating the overlap of differentially expressed genes across all compound treatments.

### Transcriptional response to αSyn aggregate formation in vitro

Modulators of αSyn primarily aim to mitigate αSyn pathology. To understand how they affect α-syn aggregation, we first assessed if and how αSyn aggregate formation itself affects transcriptional programs. We performed scRNA-seq and detected a total of 468 DEGs (**Fig. 4a**). Ontology analysis of the DEGs identified significant enrichment of protein folding and chaperone pathways, reflecting the heightened protein misfolding and aggregation induced by PFFs (**Fig. 4b**). To assess whether the DEGs were previously linked to PD, we retrieved a list of 6,253 unique genes from the Open Targets Platform^32^ that have a prior association with PD. From our DEG, 231 genes (49%) have a known PD association. These PD associated genes were significantly enriched for biological processes associated with protein misfolding and cellular stress responses (**Fig. 4d**). Molecular function analysis revealed enrichment in ubiquitination and heat shock protein binding (**Fig. 4e**). One of these PD-related genes is *CHCHD2,* and is upregulated, whereas *PRKN, UCHL-1* and *SIPA1L2* are significantly downregulated. *CHCHD2* is crucial for mitochondrial function and cellular energy production, and it is involved in the cellular stress responses, particularly in the regulation of calcium and oxidative stress^33^. *PRKN* is a key gene in maintaining mitochondrial health. It encodes an E3 ubiquitin ligase that tags damaged mitochondria for degradation through mitophagy, a critical process for cellular survival^34^. Dysfunction of *PRKN* leads to mitochondrial damage and is a well-established risk factor for PD^34^. *UCHL-1* is involved in the ubiquitin-proteasome system and dysfunction in this pathway can lead to the accumulation of misfolded proteins, such as αSyn^35^. *SIPA1L2* is known for its role in synaptic health and chromatin remodeling, also in dopaminergic neurons^36,37^. These PD-related genes are mainly involved in protein quality control, mitochondrial maintenance, and the cellular stress response, all of which are key processes implicated in the development and progression of PD. Overall, our findings indicate that transduction with PFFs induced cellular stress response pathways, which aggravate inflammation and promote cell death,^38,39^ potentially mirroring the neuronal cell death induced by αSyn aggregation. As such we see upregulation of genes involved in the unfolded protein response (UPR), a key pathway for proteostasis maintenance under cellular stress that was previously associated with αSyn accumulation in postmortem brain tissue of PD patients.^40^ For example, *HSPA1A* was upregulated and encodes a stress-induced chaperone protein involved in protein refolding mechanisms and proteotoxic stress mitigation.^41^ In addition, we observed an upregulation of genes involved in proteolytic degradation, for example *HSPA1B* and *SH3RF1*. Amongst these genes is also *SERPIN-1*, which is a negative regulator of plasminogen, an enzyme that degrades αSyn^42^. It has been proposed that increased expression of Serpin-1 can contribute to the aggregation of αSyn via impaired proteostasis^42^. Serpin-1 is also upregulated in the substantia nigra of PD patients^43^. These proteins play critical roles in protein homeostasis and managing cellular stress, especially during αSyn seeding^44^. We also observed activation of metabolic pathways that drive rRNA processing, in line with the upregulated ribosomal biogenesis accompanying αSyn overexpression in primary mouse embryonic fibroblasts.^45^ GSEA revealed activation of several cell cycle related processes, including DNA replication and chromosome segregation. Consistent with these findings, analysis of cell cycle phase distribution showed that PFF stimulation significantly reduced the fraction of cells in the G0/G1 stage, accompanied by an increase of S-phase cells (**Fig. S3d**). Finally, we observed a reduced expression of genes encoding cilium assembly and organization, in line with previous studies suggesting ciliary dysfunction induced by αSyn aggregation.^46^ Interestingly, several of these gene signatures have also been identified in postmortem PD brain tissue^47^. This further supports the use of the *in vitro* system as a PD model.

**Figure 4.**
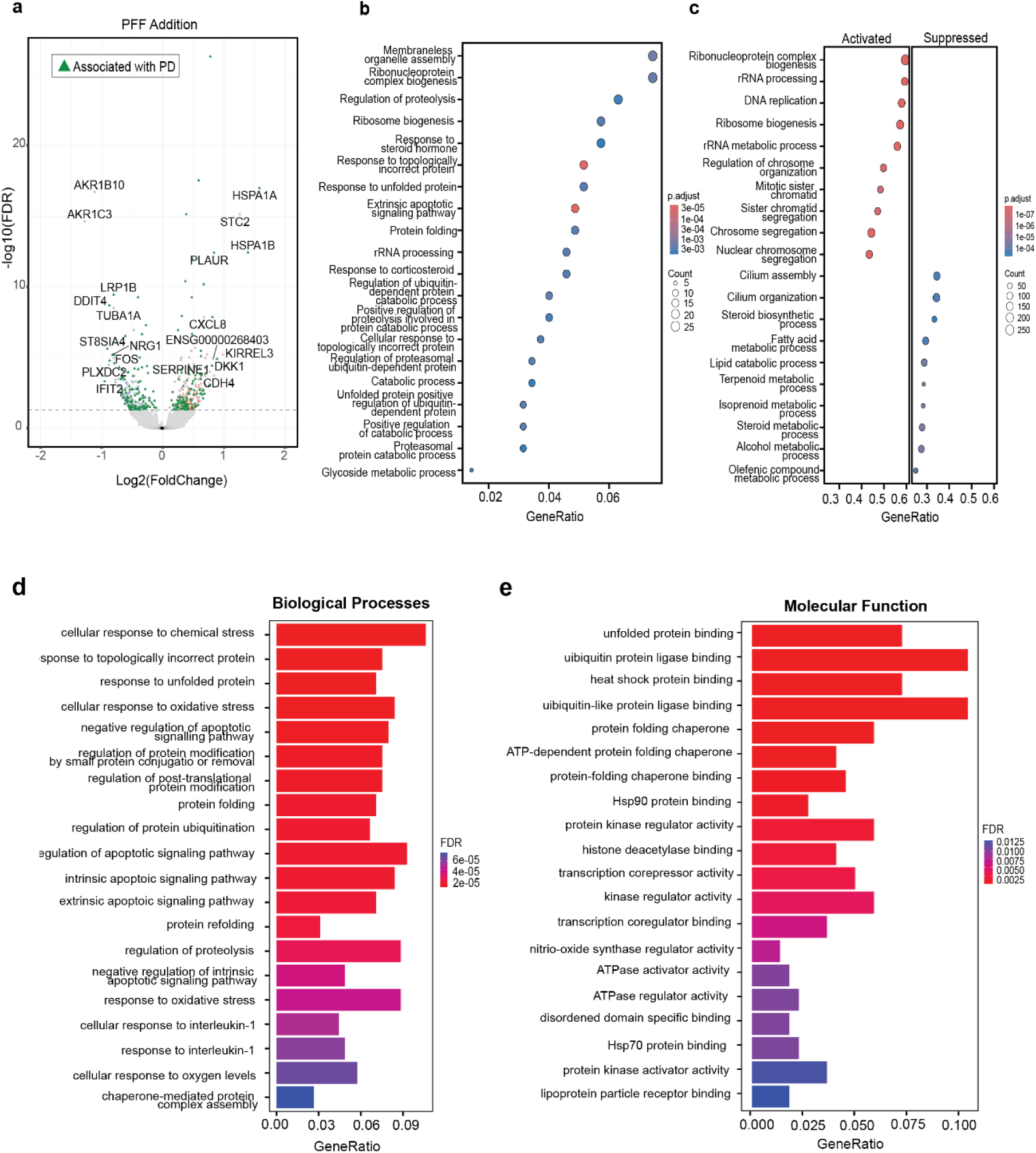
*In vitro* α-synuclein aggregation model recapitulates transcriptional signatures of Parkinson’s disease. ***(a)*** Volcano plot of DEG between cells stimulated with PFF and control cells. Each dot represents a gene; red and blue indicate significantly upregulated and downregulated genes, respectively. Green triangle indicates genes with known association with PD as indicated by Open Targets Platform. Dashed lines denote FDR = 0.05. ***(b)*** Dot plot showing the top 20 enriched biological processes (BP) identified by ORA among significantly differentially expressed genes (padj < 0.05). Dot size reflects gene count per pathway; color indicates adjusted p-value. ***(c)*** GSEA results based on all genes ranked by log₂ fold change, highlighting pathways enriched among upregulated and downregulated genes. Pathways are filtered by gene set size (100–500 genes) and significance threshold (padj < 0.05). Top 10 upregulated and 10 downregulated pathways are shown. (**d-e**) Barplots showing GO over-representation analysis using the significantly differentially expressed PD associated genes. Top 20 biological processes (e) enriched among PD associated genes, highlighting pathways related to unfolded protein response. Top 20 Molecular Functions associated with PD associated genes (d), indicating enriched of genes associated with the ubiquitination process. Color indicates adjusted p-value. Barplot height represents fraction of input genes associated with shown processes.

### αSyn modulators reverses the transcriptomic effects induced by αSyn aggregation

Having characterized the transcriptional programs induced by αSyn aggregation, we next sought to assess if and how αSyn aggregation inhibitors alter these changes. We first assessed if and how these compounds reverse the transcriptomic effects induced by αSyn aggregation, by quantifying how expression of genes in cells induced by PFF responded to treatment with each compound versus vehicle-only control (DMSO) condition (**Fig. 5a**). To further assess transcriptomic reversal, we compared log₂ fold changes of genes significantly differentially expressed in either compound +PFF versus compound alone, or vehicle-only control +PFF versus αSyn-YFP overexpressing cells. The resulting scatterplots illustrate the extent to which each compound counteracts PFF-induced gene expression changes (**Fig. 5b**). Interestingly, out of 468 genes showing differential expression upon PFF transfection, 411, 430 and 465 were no longer differentially expressed upon Minzasolmin, Emrusolmin and EGCG treatment, respectively (**Fig. 5c**). Overall, these observations support the notion that these compounds reduce αSyn aggregation and the ensuant transcriptional responses in this *in vitro* PD model. Gene ontology analysis of this group revealed enrichment in biological processes related to rRNA and protein localization processes (**Fig. 5d**). Notably, several pathways pointed to the involvement of the nuclear Cajal body organelle, which plays a critical role in regulating ribosome biogenesis, small nuclear ribonucleoprotein (snRNP) maturation, and telomere maintenance. In addition, significantly altered processes such as membraneless organelle assembly may also contribute to the formation of other membraneless organelles, including processing bodies (P-bodies), which function in mRNA turnover and storage. Previous studies have shown that αSyn can interact with core components of these RNA regulatory complexes, including EDC4, a key scaffold protein of P-bodies.^48^

**Figure 5.**
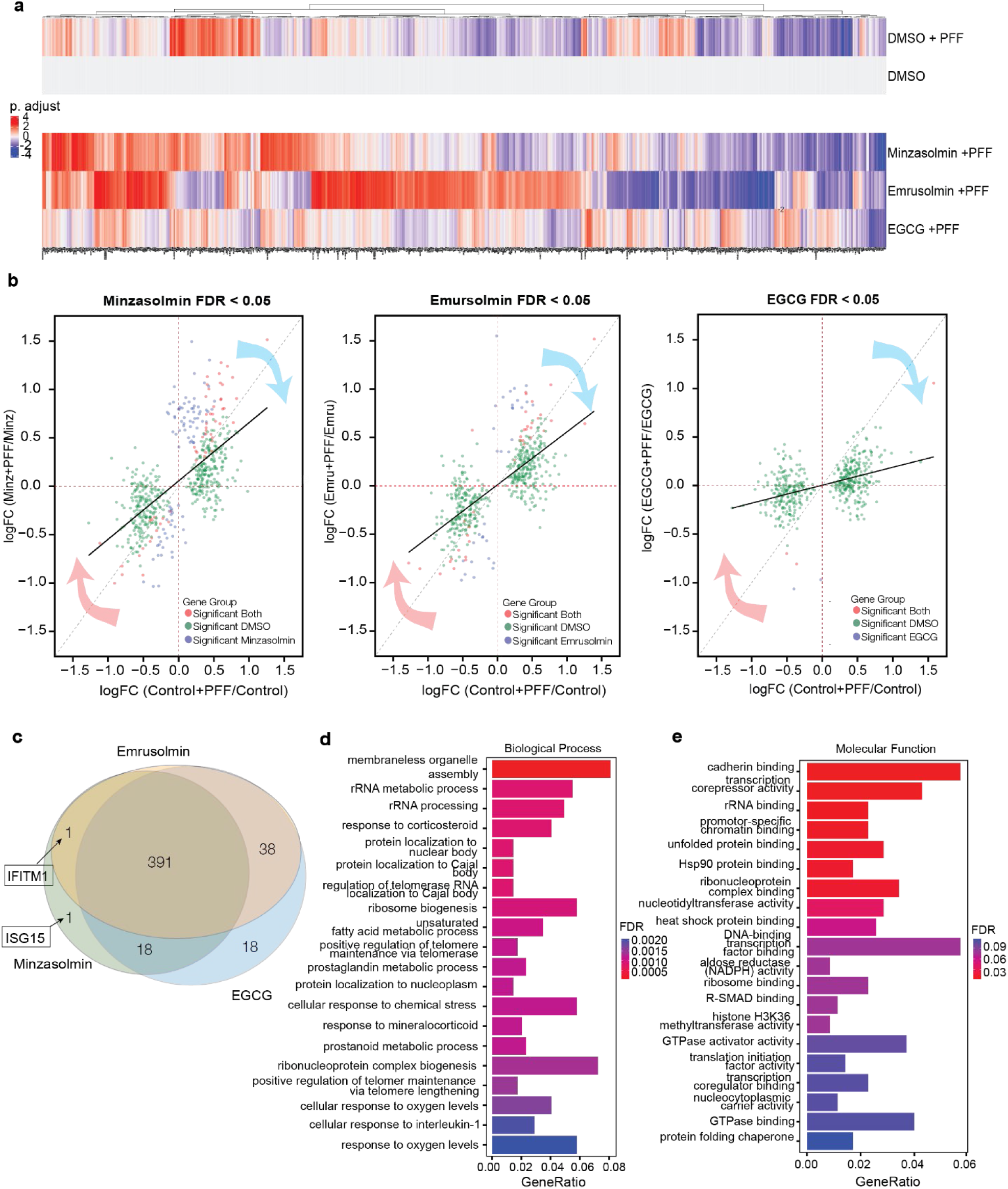
Transcriptomic reversal of PFF-induced transcriptional signatures by α-synuclein aggregation inhibitors. ***(a)*** Heatmap of differentially expressed genes across all conditions compared to DMSO-treated controls, colored by adjusted p-value. ***(b)*** Scatterplots showing genes with FDR < 0.05 in either comparison. The x-axis displays log₂ fold changes between DMSO + PFF versus DMSO alone condition; the y-axis shows log₂ fold changes between compound + PFF and compound-only condition for Minzasolmin (left), Emrusolmin (middle), and EGCG (right). Each point represents one gene. The diagonal dotted line shows X = Y, demonstrating the trend if effect sizes were equal, black solid line is trendline of the observed transcriptional signatures. Red and blue arrows indicate the direction of reversal of the PFF induced effect. ***(c)*** Venn diagram illustrating the overlap of genes significantly upregulated in the DMSO + PFF condition that are no longer differentially expressed following compound treatment for all three compounds, Minzasolmin, Emrusolmin and EGCG. (**d-e**) Bar plots of GO over-representation analysis for the 391 reversed genes. Bar color indicates adjusted p-value. Shown are the top 20 enriched biological processes (d) and top 20 molecular functions (e). X-axis is showing GeneRatio of reversed gene set in enriched terms.

For molecular function ontology, enrichment in this gene set was most pronounced for transcription corepressor activity, transcription coregulator activity and cadherin binding (**Fig. 5e**). These terms highlight a link between the reversed genes and transcription regulation. Together, this indicates a central role for the pathology associated genes in nuclear organization, RNA processing and overall genome stability.

### Inhibitor-specific gene expression signatures highlight unique mode of actions

Lastly, we assessed the effect of αSyn modulators on the cells transcriptome in the PFF induced, pathological condition. Treatment with Minzasolmin, Emrusolmin, and EGCG in PFF exposed cells resulted in 137, 638 and 136 DEGs (**Fig. 6a-c**), relative to the PFF exposed DMSO control. Only six genes were commonly regulated across all three αSyn aggregation inhibitors (**Fig. 6d**). Gene ontology enrichment analysis of this shared subset revealed biological processes related to RNA metabolism and ribosome function, many of which were consistently attenuated across conditions. In contrast, genes involved in biosynthesis and hormone metabolic pathways were upregulated, suggesting a shift in cellular energy allocation away from ribosomal activity toward essential metabolic functions. This shift may reflect a direct effect of compound activity in mitigating α-synuclein–induced disruption, or alternatively, an indirect consequence of reduced aggregate formation. Together, this suggests a shared transcriptional response to α-synuclein aggregation. This convergence on similar transcriptional pathways may reflect a compensatory mechanism aimed at restoring cellular homeostasis under proteotoxic stress.

**Figure 6.**
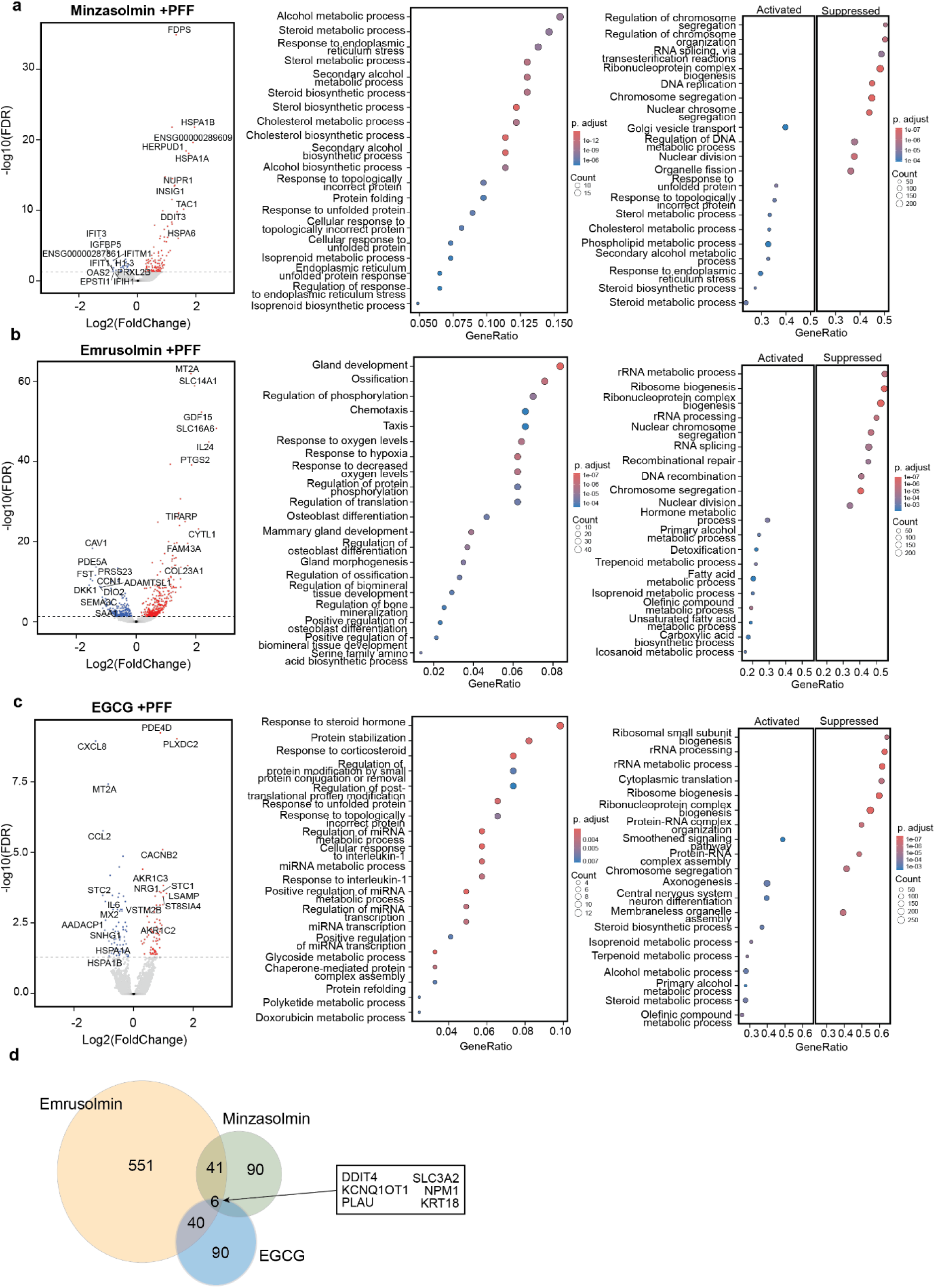
Differential gene expression analysis reveals compound-specific modulation of PFF-induced signatures. ***(a-c)***. Volcano plots (Left) showing differential gene expression between PFF stimulated cells treated with Minzasolmin (a), Emrusolmin (b), or EGCG (c) and DMSO +PFF-treated controls. Each dot represents a gene; red and blue indicate significantly upregulated and downregulated genes, respectively. Dashed lines denote FDR = 0.05. Dot plot showing the top 20 enriched biological processes (BP, middle) identified by ORA among significantly differentially expressed genes (padj < 0.05). Dot size reflects gene count per pathway; color indicates adjusted p-value. Dot plots displaying gene set enrichment analysis (GSEA, Right) results based on all genes ranked by log₂ fold change, highlighting pathways enriched among upregulated and downregulated genes. Pathways are filtered by gene set size (100–500 genes) and significance threshold (padj < 0.05). Top 10 upregulated and 10 downregulated pathways are shown. ***(d)***. Venn diagram illustrating the overlap of differentially expressed genes across all compound treatments in PFF induced states. The 6 overlapping differentially expressed genes are annotated.

In addition to the pathways involved in RNA processing and core metabolic functions, which we saw as consistently normalized by all three compounds, several additional, biologically significant pathways were enriched. These included processes uniquely modulated by individual compounds, highlighting both shared and compound-specific mechanisms of action. Notably, several cell cycle related processes, including chromosome segregation, were enriched following treatment with the inhibitors. This effect was particularly pronounced for Minzasolmin, where approximately half of the suppressed pathways were associated with cell cycle progression, such as DNA replication and chromosome segregation. However, analysis of cell cycle phase distribution revealed that Minzasolmin, both in the presence and absence of PFF, did not significantly alter the proportion of cells in the G1 phase compared to the DMSO control (**Fig. S3d**). This suggests that Minzasolmin addition effectively reversed the PFF-induced activation of cell cycle pathways, which were elevated in the DMSO +PFF condition versus vehicle-only (DMSO) control and accompanied by a measurable shift in cell cycle proportions. On the other hand the cell cycle distribution was still significantly altered in the EGCG +PFF condition compared to control. Both Emrusolmin and EGCG treatment induced genes involved in overlapping metabolic pathways, including olefinic compound metabolism, isoprenoid biosynthesis, and cellular hormone-associated metabolic processes. GO analysis revealed that Emrusolmin modulates cellular responses to oxygen concentration changes, with significant enrichment of genes involved in the hypoxia response pathway, which may be relevant given the established link between oxidative stress and dopaminergic neuronal loss in PD.^49^ Interestingly, functional enrichment analysis of DEG in the Emrusolmin-treated condition revealed several pathways not typically associated with neurodegenerative contexts, including ossification, and (mammary) gland development. These findings may point to previously unrecognized off-target transcriptional effects of Emrusolmin or reflect broader transcriptional shifts induced by the compound.

Several gene signatures observed in compound-treated cells under pathological conditions were also previously enriched in the non PFF induced cells exposed to the same inhibitors. This suggests that these transcriptional responses are primarily driven by the compounds themselves and may reflect their unique modes of action, rather than being secondary effects of αSyn aggregation reversal. These responses could also indicate potential off-target effects unrelated to αSyn disaggregation. For instance, Emrusolmin modulated genes involved in RNA splicing and xenobiotic metabolism (e.g., *MT2A*), Minzasolmin selectively induced stress response genes (e.g., *HSPA1A*), and EGCG enriched TNF-related genes, with no pathways significantly enriched in GSEA without PFF. More pathways were enriched in PFF-treated versus control conditions, underscoring the context-dependent nature of compound activity, primarily yielding transcriptional responses in the pathology model. The most pronounced transcriptomic effects of the compounds were PFF-dependent, such as the suppression of rRNA processing following PFF exposure. In summary, all three inhibitors effectively reduced αSyn aggregation in the cellular PFF seeding model. Transcriptionally, they counteracted PFF-induced perturbations through both shared and compound-specific mechanisms.

## Discussion

This study presents a robust and scalable cellular αSyn seeding assay to screen small molecules for drug discovery purposes. The cellular seeding assay mimics the pathogenic aggregation of human αSyn in a controlled cellular environment, allowing for medium to high-throughput screening and detailed phenotypic analysis. By using αSyn-YFP as a fluorescent reporter, it is possible to quantify seeded aggregation in a non-biased and automated way.

The addition of recombinant human PFFs to αSyn-YFP or αSyn overexpressing cells initiates a robust seeding process that drives αSyn aggregation, and which resembles key pathological features observed in synucleinopathies^1,3^. The observed colocalization of αSyn aggregates with PSer129-αSyn and other aggregation-based markers further validates the presence of pathological inclusions. The scalability of the assay between different well formats (24, 96 and 384 well plates) allow its application in both phenotypic screening and transcriptomic profiling, bridging the gap between cellular models, molecular pathway analysis and drug discovery.

The seeding with fibrillar aggregates causes significant transcriptional changes of cellular pathways with strong activation of protein homeostasis mechanisms, including the UPR and chaperone-mediated protein folding pathways. Additionally, pathways involved in autophagy and proteolysis are activated, underscoring the cellular effort to degrade aggregated proteins and maintain proteostasis. Lipid metabolism pathways are also significantly altered, likely due to the interaction of αSyn with lipid membranes, as the cellular membrane is an important catalyst of αSyn aggregation^50–52^. Together, these activated pathways highlight the cellular stress responses that are more aberrantly triggered by αSyn fibrillar seeding.

We tested three molecules with known and documented inhibitory effects on αSyn aggregation^22,24,53^. All three compounds effectively inhibited αSyn aggregation, and transcriptomic analysis revealed consistent enrichment in metabolic and rRNA-related processes. The effect on lipid metabolism was particularly relevant, given its role in αSyn aggregation and the role of well known and documented PD risk genes such as *GBA1* which encodes for the glucocerebrosidase enzyme^54,55^. Mutations in this gene are one of the most important genetic risk factors for the development of PD. High molecular weight assemblies of αSyn are known to closely interact with lipid membranes, influencing synaptic function and promoting ROS production^56–59^. Inhibitor-induced disaggregation may reduce ROS, restoring metabolic activity and relieving stress responses such as the unfolded protein response (UPR). Furthermore, lipid composition impacts α-syn aggregation in PD models^60,61^.

EGCG can directly bind to aggregated and non-aggregated forms of αSyn^19,20^. EGCG was first extracted as the major phenolic compound in green tea. It proposedly modulates α-syn pathology either by binding αSyn monomers and diverting oligomerization from toxic towards non-toxic forms ^62^, or by remodeling large αSyn fibril aggregates into smaller, non-toxic aggregates^63^, preventing neurotoxicity.^64^ EGCG also inhibits aggregation of other amyloid proteins, suggesting a broad potential to treat neurodegenerative diseases.^65^ In the cellular assay EGCG showed a protective role, enhancing protein homeostasis and mitigating stress-related pathways. The effects of EGCG are limited in the overexpression assay, with only 9 DEGs whereas EGCG has a much stronger impact in the seeded aggregation assay. These gene expression changes reflect pathways related to ribosome biogenesis and inflammation-related pathways. They highlight the protective role of EGCG for multiple pathomechanisms, particularly by ameliorating the heightened stress caused by seeded αSyn aggregation. Due to its strong anti-inflammatory and anti-oxidant effects, EGCG has been evaluated for use in various pathologies, from colorectal cancer to HIV.^66–68^ Although its potential beneficial effects have been shown in preclinical work, EGCG failed to show clinical benefits in some advanced clinical studies in PD^53,69^. EGCG was also tested in a phase 2/3 clinical study in MSA, but it did not meet the study endpoints^69^. There was also a higher incidence of hepatic adverse events in the EGCG group at higher doses^69^. Even in the light of the potential therapeutic effects of EGCG it has therefore been questioned if EGCG or other polyphenols are good treatment options for synucleinopathies ^70^.

Emrusolmin is a small molecule that directly binds aggregated αSyn^18,71^. Emrusolmin is an aggregation inhibitor that binds αSyn fibril in a structure-dependent manner. It inhibits aggregate formation, blocking development of pathological aggregates.^72^ Emrusolmin treatment prolonged survival of prion-infected mice by inhibiting aggregate formation^73^, and also alleviated the symptoms of amyloid beta pathology in a mouse model of Alzheimer’s disease.^74^ It has been proposed as a scaffold for PET imaging and as a potential therapeutic molecule for synucleinopathies. In the αSyn-YFP overexpression and seeding assays, Emrusolmin exhibited strong effects on cellular structural responses and oxidative stress pathways, including upregulation of genes such as *MT2A* and *PTGS2*. While these changes underscore its therapeutic promise, Emrusolmin also positively regulates apoptotic pathways in the cellular seeding pathway, and additional research is warranted to understand the contribution of these pathways. Nevertheless, Emrusolmin has shown promising effects in animal models of PD and MSA showing a reduction of αSyn pathology and a reduction of cellular loss ^75,76^. Emrusolmin is currently in early clinical development for the treatment of PD NCT04685265^23^. A phase 2 clinical trial is planned for people with MSA to assess safety and efficacy (NCT06568237)^77^.

Minzasolmin is a small molecule designed to inhibit αSyn multimerization. Minzasolmin is an orally available small molecule that increases the flexibility of membrane-bound αSyn, thus weakening its membrane association. Multimerization of αSyn happens naturally at the membrane surface^50,51^ and Minzasolmin has been shown to bind with αSyn multimers at the membrane interface after chemical cross-linking^17^. It promotes release of αSyn monomers into the cytosol to prevent toxic fibril formation.^78^ In the aggregation assays Minzasolmin demonstrated a significant impact on lipid metabolism and protein folding pathways, including the upregulation of *FDPS* and *INSIG1*, which could be linked to membrane composition and αSyn multimerization or aggregation dynamics. The suppression of *IFIT* genes indicates a dampening of pro-inflammatory and antiviral responses. IFIT and IFITM proteins are activated by type-1 interferon and their collective suppression could indicate a dampening of inflammatory processes and reduced activation of antiviral and immune responses^79^. Given the role of inflammatory or infectious triggers in PD and MSA^80,81^, an anti-inflammatory role of this therapeutic agent could be useful, especially during early stages of disease, or in patient groups that are specifically susceptible for infections or inflammation. Minzasolmin has been shown to ameliorate αSyn pathology in preclinical models^82^ and has been shown safe in phase 1 clinical studies^83^. It has progressed to a phase 2 clinical trial (ORCHESTRA, NCT05543252, NCT04875962 and NCT04658186) to test efficacy, safety and pharmacokinetics in people with PD. As of 2025 the last study was terminated due to failure to meet the study endpoints.^84–86^

The actions of these three compounds are also highlighted by the activation or suppression of unique pathways, with sometimes opposing effects. For instance, EGCG suppresses heat shock protein activity, potentially mitigating the protein-folding stress caused by aggregation. This could be linked to its direct binding with fibrils, or to its broader anti-inflammatory effects and its role in stabilizing cellular environments under stress. EGCG its selective impact might indicate a pathway favoring reduced activation of heat shock responses, which could limit unnecessary cellular stress responses during aggregation. Conversely, Minzasolmin, that also directly binds αSyn, upregulates heat shock proteins (*HSPA1A* and *HSPA2A)*, suggesting it actively promotes cellular mechanisms to manage protein misfolding and aggregation. The upregulation of these proteins indicates a more direct engagement with proteostasis pathways, stimulating cellular capacity to refold misfolded proteins or target them for degradation. These opposing effects may reflect to some extent the differential interactions with αSyn and regulation of cellular stress mechanisms, but it also underscores the complexity of targeting αSyn aggregation.

While the assay provides a powerful platform for studying αSyn aggregation, limitations remain. The use of a single cell line, H4 neuroglioma cells, may limit the generalizability of findings to other cell types or *in vivo* conditions. The current study works to underline the possibilities of this platform and an explorative screen to study the affected transcriptional signatures. Follow up studies should incorporate additional screening tools, such as primary culture systems or humanized models, such as iPSC-derived cell models to validate findings from an initial medium or high-throughput screening model. The scalable cellular seeding assay developed in this study offers a valuable tool for dissecting αSyn aggregation and its modulation by therapeutic compounds. By integrating phenotypic and transcriptomic analyses, this work contributes to a deeper understanding of αSyn-related pathologies. Understanding these changes is crucial for identifying potential therapeutic targets, understanding disease mechanisms, or assessing the effects of drugs or other interventions. Collectively, helps to advance the search for effective interventions in PD and MSA

## Acknowledgements

We utilized OpenAI ChatGPT to assist in the proofreading and refinement of this manuscript.

## Author contributions

Conceptualization: WP, PVM, BT, CVH, VD, VB

Methodology: WP, PVM, CVH

Investigation: CVH, PVM, QY, XS, SH

Funding acquisition: WP, VD, VB

Project administration: VD

Supervision: CVH

Writing Initial draft: PVM, WP, VD

Writing – review & editing: WP, PVM, CVH, VD, BT

## Data and materials availability

All data will be available upon request.

## Supplementary Figures

**Supplementary Figure 1.**
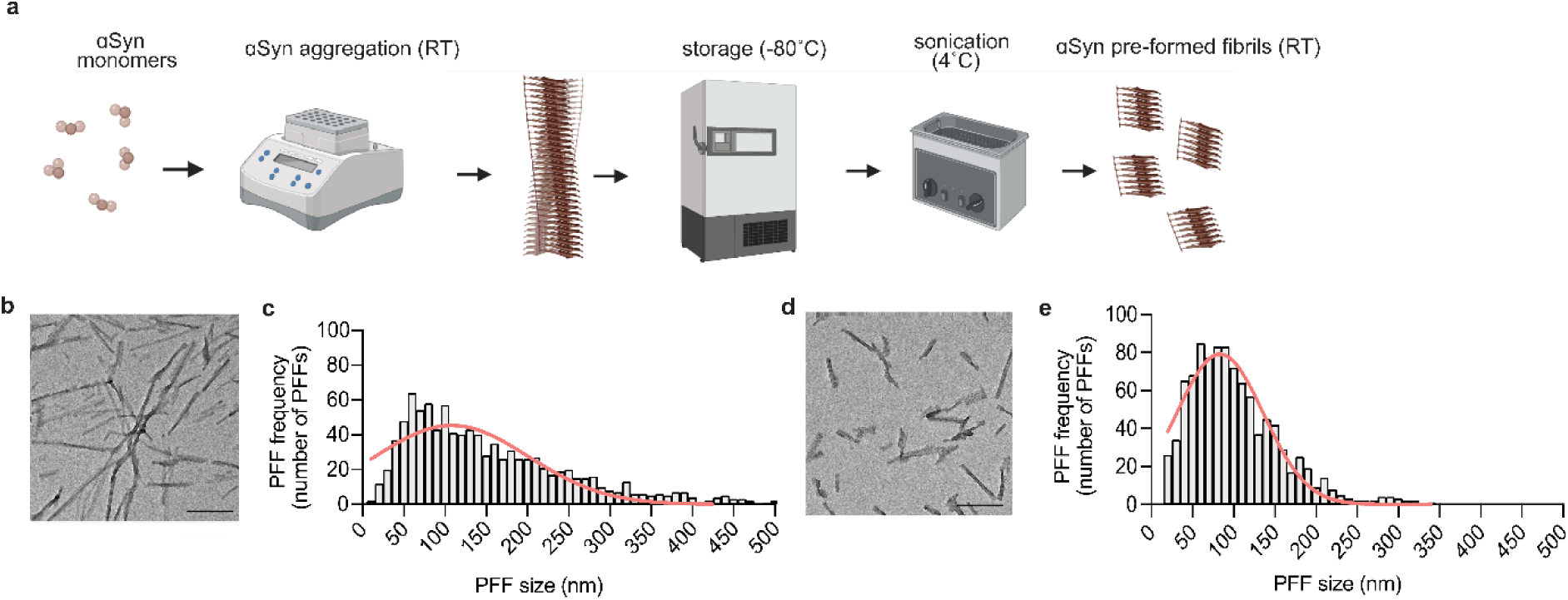
Characterization of pre-formed fibrils (PFFs). Schematic representation of the PFF aggregation process. ***(a)*** Recombinant ɑSyn monomers are aggregated *in vitro* under controlled conditions and immediately stored at −80°C for later in single use aliquots. After thawing, fibrils are sonicated in a water bath into ready to use PFFs. ***(b)*** Transmission electron microscopy (TEM) image showing long fibrils prior to sonication. Scale bar: 200 nm. ***(c)*** Size distribution of pre-sonicated fibrils, measured as PFF frequency against fibril length. ***(d)*** TEM image showing PFFs after sonication, demonstrating a reduction in fibril size. Scale bar: 200 nm. ***(e)*** Size distribution of sonicated PFFs, showing the frequency of PFF lengths with a peak around 50–100 nm. The PFF characterization confirms efficient breakdown of fibrils into smaller, more uniform aggregates suitable for cellular seeding assays.

**Supplementary Figure 2.**
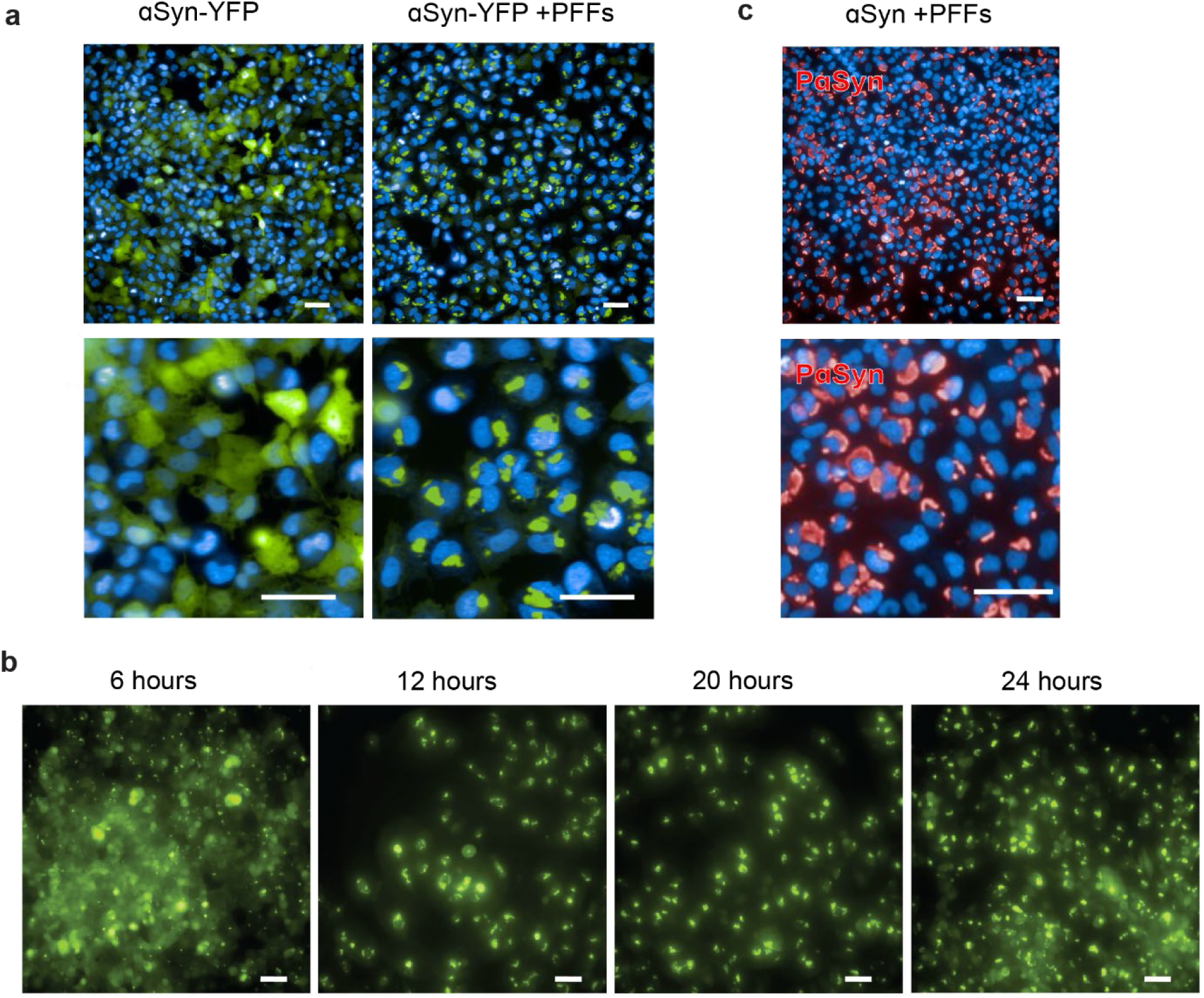
Immunofluorescence visualization of PFF-induced α-synuclein aggregation in cells expressing αSyn with or without YFP. ***(a)*** Representative images showing ɑSyn-YFP fluorescence in untreated cells (left) and cells treated with PFFs (right). Diffuse fluorescence is replaced by discrete puncta indicative of aggregation. Nuclei are stained with DAPI (blue). Scale bars: 50 µm (top) and 20 µm (bottom). ***(b)*** Time-course analysis of seeded aggregation in αSyn-YFP–overexpressing cells over 24 hours following PFF exposure. Puncta are detectable as early as 6 hours and persist through 24 hours. Scale bars: 20 µm. ***(c)*** Immunocytochemical detection of pathologically phosphorylated α-synuclein (pαSyn, red) in cells treated with PFFs. Cells overexpress αSyn without a fluorescent tag. Scale bars: 50 µm (top) and 20 µm (bottom).

**Supplementary Figure 3.**
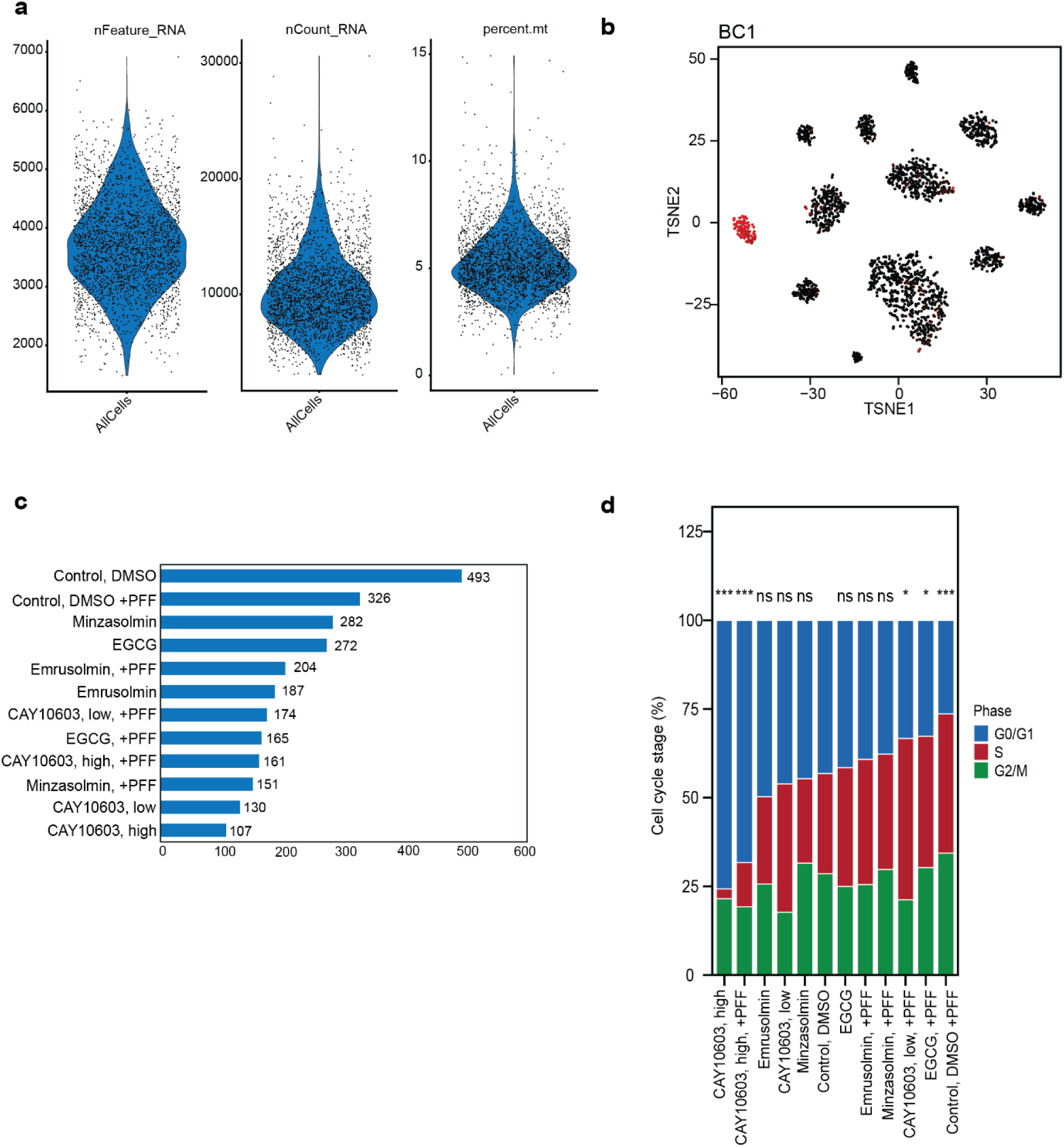
Quality control for scRNA-seq. ***(a)*** Quality control for scRNA-seq data after filtering, cells with RNA count between 1.500-20.000 and mitochondrial percentage <15% were retained. ***(b)*** t-SNE embedding of cells colored by log-transformed BC1 counts (black to red). A distinct cluster of BC1-enriched cells is visible, indicating robust barcode signal and supporting singlet assignment. ***(c)*** Barplot showing cell number for each condition within the screen ***(d)*** Barplot indicating cell cycle stage distribution across conditions, with “***” indicating p < 0.001, “**” is p < 0.01, “*” is p < 0.05 and ns = not signifcant. ***(e)*** UMAP projection of single-cell transcriptomes, separated by treatment condition and absence or presence of PFF.

**Supplementary Figure 4.**
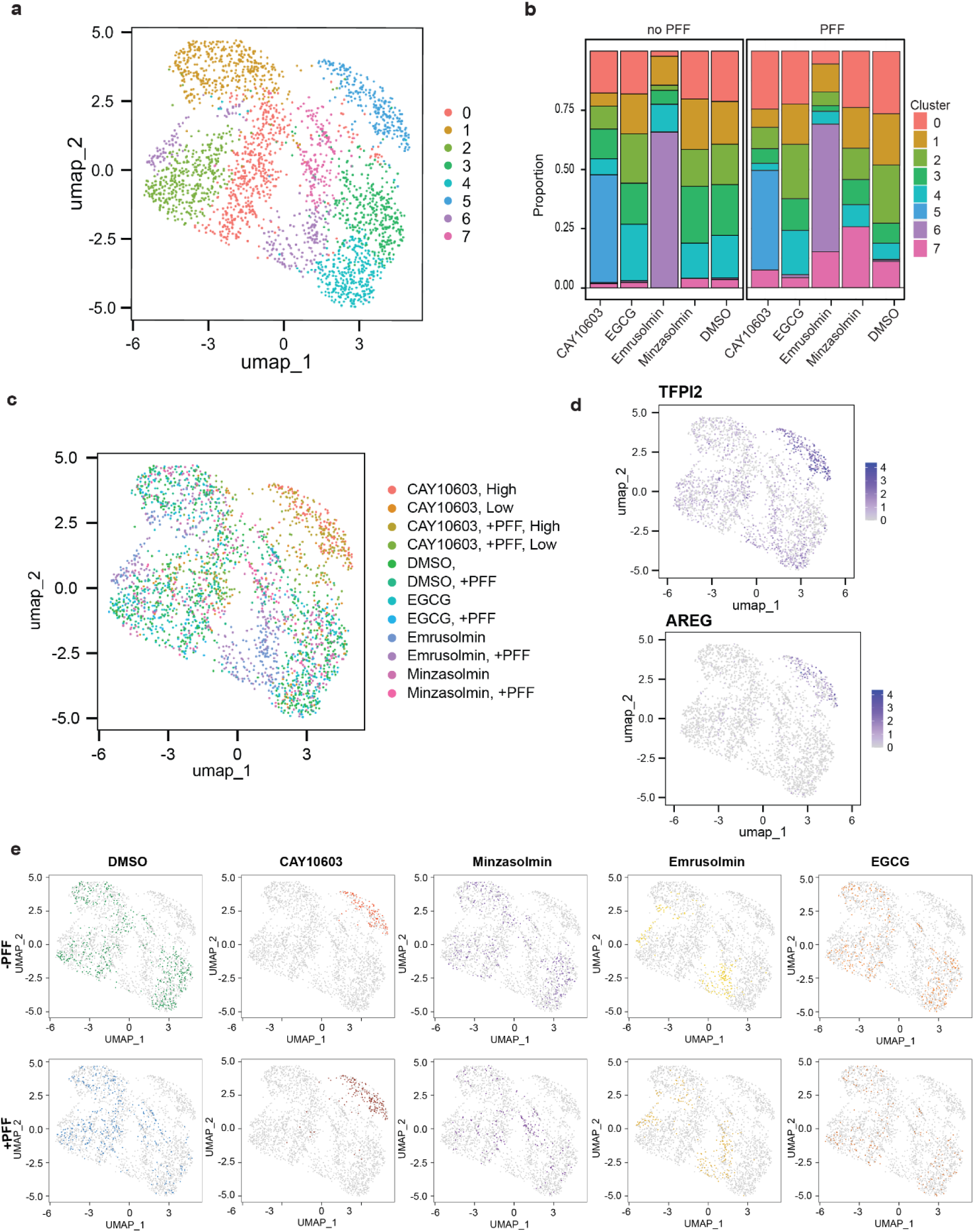
UMAP visualization of clustered cell populations across conditions. ***(a)*** UMAP projection of single-cell transcriptomes, with cells colored by Seurat-defined clusters. ***(b)*** Stacked bar chart showing the relative abundance of transcriptional clusters across compound-treated conditions under naïve and PFF-stimulated states, highlighting distinct shifts in cell state composition induced by α-synuclein aggregation and its inhibition ***(c)*** UMAP projection of single-cell transcriptomes, with cells colored by treatment condition. ***(d)*** UMAP feature plots showing expression of *AREG* and *TFPI2*, which are enriched in cluster 5. **(e)** UMAP projection of single-cell transcriptomes, separated by treatment condition and absence or presence of PFF.

